# The eukaryotic horizontal gene transfer dataset a compendium

**DOI:** 10.1101/2025.11.05.686818

**Authors:** Johnathan A. Spaulding, Janna L. Fierst

## Abstract

With more eukaryotic genomes available for study researchers have been able to identify a growing number of horizontal gene transfer (HGT) candidates. We compiled 9,511 protein coding genes that were identified as horizontally transferred in the published literature. This dataset contains gene transfers from bacteria, fungi, archaea and protists to metazoans. We assigned a level of certainty to each gene based on the methods used in the scientific paper reporting HGT. A supplemental file contains all the coding sequences and protein sequences for the HGT genes. This dataset can be used to identify trends in genome and protein evolution and provide a foundation for creating a centralized HGT database for eukaryotes.

## Introduction

### Background & Summary

Horizontal or lateral gene transfer (HGT) is the movement of genetic material (DNA) from one organism to another, separate from parent to offspring (1). This mode of genetic transfer was first discovered in bacteria and accounts for a significant amount of microbial genetic diversity (2). Interestingly, the genes that are transferred to recipient bacteria often perform biological functions that give the recipient an evolutionary advantage (3). For instance, bacteria that are susceptible to antibiotics can acquire antibiotic resistance genes via HGT from antibiotic resistant bacteria in the area (2). This leads to an expansion in the ecological niche for the newly antibiotic resistant bacteria, illustrating the importance of HGT.

In contrast to prokaryotes, HGT is rare in eukaryotes (4). The scarcity of verified HGTs in eukaryotes could be due to technological or biological reasons. Technologically, there could be sequencing bias, in which organisms that are typically sequenced aren’t the organisms receiving HGTs (5). Biologically, eukaryotic organisms may be less prone to acquire HGTs due to the physical separation of germ-line cells from somatic cells. Somatic cells may shield the germline from foreign DNA, decreasing the chance of foreign DNA being transmitted to the progeny (6).

While eukaryotic organisms may not engage in HGT as frequently as prokaryotes, what they do have in common is that transferred genes can provide adaptive functions potentially aiding the recipient. For example, there have been multiple cases of plant parasitic nematodes gaining foreign genes from bacteria and fungi that facilitate parasitism (7). The plant parasitic nematode Bursaphelenchus xylophilus has gained multiple genes from fungi that have aided it in parasitism such as plant cell wall degrading enzymes (8). To identify more cases of eukaryotic HGT and study its biological significance, more eukaryotes from neglected groups need to have their genomes (5) and transcriptomes (9) sequenced and analyzed.

Advances in machine learning and statistical methods could form the foundation for rigorous bioinformatic identification of HGTs but methods development has been slowed by a lack of validated HGTs (10). In the absence of a comprehensive empirical dataset researchers developing HGT identification methods have used simulated datasets to simulated HGT events by manipulating the biological sequences. This frequently involves creating synthetic chimeric genomes by inserting known foreign DNA sequences into the recipient genome at random, with the goal of testing a novel method to identify HGT in genomes (10,11). For example, Jaron et al simulated HGT by randomly selecting and replacing DNA sequences across 10 genomes, then applied their new method, SigHunt to identify the inserted sequences (11). Simulations may approximate very recent HGTs but over time these sequences will be subject to mutation and selection, changing the foreign origin ‘signal’ HGT identification methods rely on.

Fortunately, biology is experiencing an unprecedented increase in the amount of data being generated (12,13). The cost of sequencing genomes has decreased for the last two decades (14). This, in turn, has enabled more laboratories to generate rich genomic datasets (15). In the current literature HGTs are increasingly being identified across diverse organisms but no comprehensive dataset exists to collect HGTs published in past and present papers. A major bottleneck in studying HGTs stems from the fact that this information, although growing, remains confined to individual publications.

In this study, we created a dataset that contains 9,511 HGT candidates identified in published articles between 2000 and 2025. This dataset contains HGTs from bacteria, fungi, archaea, and protists to metazoan recipients. We also provide both tested and untested predictor variables for the transferred genes which may be used to statistically differentiate HGTs from native sequences. Our tested predictor is Guanine-Cytosine (GC) content (16), while our untested predictors are amino acid length and coding sequence length. This dataset will allow researchers to comprehensively analyze sequence data, discover emergent patterns of HGTs and provide a foundation for creating a centralized HGT database for eukaryotes.

## Methods

### Workflow for eukaryotic HGT dataset (euHGT) paper search

To create the euHGT dataset, we identified articles using a Web scraper program called ScrapPaper that extracts information from Pubmed search results (17).The program extracted the titles of all research papers in the published literature containing the keywords “ Lateral Gene Transfer”, “Horizontal Gene Transfer”, and “metazoan”(https://pubmed.ncbi.nlm.nih.gov/?term=Lateral+Gene+Transfer%2C+Horizontal+Gene+Transfer%2Cmetazoan&filter=years.2000-2025) from 2000 to 2025. The output of this program displayed a dataset containing titles of research papers identifying prokaryote/fungi/archaea to eukaryote transfers, prokaryote to prokaryote transfers, and viral to eukaryote transfers. Only the research papers pertaining to bacteria-, fungi-,archaea-, and protist-to-eukaryote transfers were used to create the euHGT dataset. Thus, out of the 151 HGT papers that were identified using ScrapPaper only 36 papers were included in the euHGT dataset.

For each article the transferred nucleotide gene sequence and corresponding amino acid sequence were obtained from the National Center for Biotechnology Information (NCBI) or the data deposition website or database reported by the authors. The transferred gene accession number, genome assembly ID, protein accession number, gene name/function, phylogenetic support, level of certainty, inferred donor taxon (Bacteria, Fungi, Protist, or Archaea), host taxon, and paper reporting the HGT(s) were also recorded when possible. Custom Python scripts, available at https://github.com/Genome-explorer were used to calculate the Guanine-Cytosine percentage, gene length, protein length, and coding sequence length for each of the HGTs. An outline of our workflow is displayed in Figure 1.

**Figure 1.**
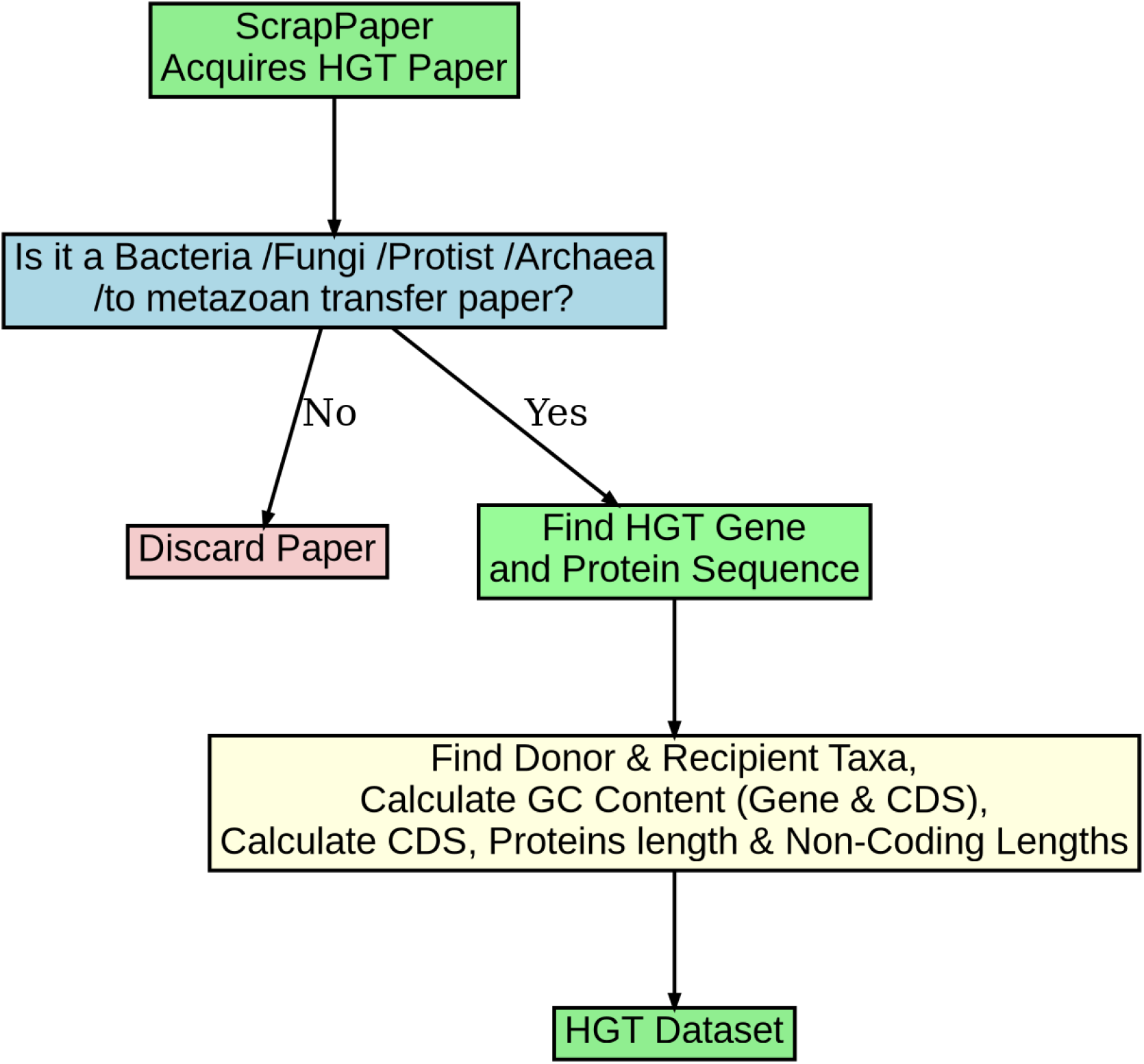
Data collection workflow for the euHGT dataset. Each paper was selected based the type of HGT. The euHGT dataset focuses on bacteria-, fungi-, archaea-, and protist-to-eukaryote transfers. Once the paper was selected the next step was to locate the gene and protein sequences.

### Certainty metric

We created a certainty metric to assess the likelihood of the reported gene transfer being a true HGT. Both errors in sequencing and contamination can lead to false positive HGT identification. Experimentally, multiple steps in the wet lab including library construction, amplification of DNA and sequencing can introduce errors (17). Downstream effects of these processes can lead to gene prediction errors including false sequences that are not present in the organism (18). The end result of this scenario mirrors contamination, with DNA from a foreign organism being mixed with the primary organism being sequenced. A few studies have shown previously identified HGT genes as contamination (19,20,21).

To increase confidence that a gene is a true HGT, multiple methods should be used to assess both presence in the genome and evolutionary origin (22). Thus, our certainty metric has five levels with higher levels signifying a greater confidence in the gene transfer event (Table 1). Each gene is assigned a certainty score based on multiple lines of evidence provided in the original paper reporting the transfer event. Both empirical laboratory evidence and computational analyses were weighted depending on reliability of the techniques and potential biological information. For example, papers that included the use of both in situ hybridization (or RNAi or Crispr) and phylogenetic analysis have a level five certainty. In situ hybridization is a molecular technique that shows the location of a specific sequence of DNA in the biological sample, which excludes the chance that the gene is a result of contamination (23). Phylogenetic analysis is a reliable statistical method for identifying evolutionary gene origin and confirming sequences have been horizontally transferred (24).

**Table 1:** Levels of certainty and the required conditions for each level. Higher levels of certainty signify a greater confidence in the gene being the result of a true horizontal transfer.

| Level Of Certainty | Criteria | Evidence Description |
| --- | --- | --- |
| 5 | In situ hybridization +<br>Phylogenetic analysis | Confirms gene location, excludes contamination; |
|  |  | phylogeny supports HGT origin |
| 4 | RNA validation + Phylogenetic analysis | Gene is expressed; phylogenetic analysis supports evolutionary history |
| 3 | PCR or Southern blot + Phylogenetic or sequence-based analysis | Confirms presence (with contamination risk); phylogeny or sequence adds reliability. |
| 2 | Sequence-based method + Phylogenetic analysis | Sequence similarity + phylogeny |
| 1 | Only one method (e.g., sequence or phylogeny) | Only one line of evidence is used. |

Studies that utilized both empirical RNA-based methods including quantitative PCR, ESTs, or the Race method and phylogenetic or sequence-based analysis were assigned a level four certainty. RNA evidence demonstrates that the gene is actively expressed in the organism (25). Research papers using both empirical DNA-based methods including Polymerase Chain Reaction (PCR), southern blotting or DNA probes (26) and phylogenetic methods or sequence-base analysis (27) have a certainty level of three. Using DNA-based methods can confirm the presence of the gene in the biological sample potentially amplify intra- and inter-cellular contamination (28). Papers that used both a sequence-based method (27) such as BLAST (Basic Local Alignment Search Tool) and phylogenetic analysis were assigned a level two certainty. Lastly, papers that only use one method of detection such as sequence-based methods or phylogenetic analysis or empirical DNA-based methods or RNA-based methods were assigned a level one certainty.

### Recurrent top 7 phyla that experienced HGT

After collecting transferred genes present in different organisms, we investigated patterns in HGT. The top seven phyla that experienced the most gene transfers were: Rotifera, Arthopoda, Nematoda, Chordata, Porifera, Echinodermata, and Mollusca. This was calculated by the number of transferred genes present in each species that belong to their respected phyla. Phylum Rotifera had the highest number of transferred genes out of all the phyla present in the dataset (Figure 2). The phyla with highest number of transferred genes with unknown origin were Nematoda (298) and Porifera (190; note-, the log scale makes the unknown bar in Porifera appear larger than the Nematoda unknown bar). These transferred genes had a non-metazoan origin reported in the paper but the authors also reported uncertainty in assigning the donor organism. Phylum Mollusca received the fewest HGTs of these 7 phyla (Figure 2). Overall, each of the top 7 phyla had the most foreign genes originating from bacteria.

**Figure 2.**
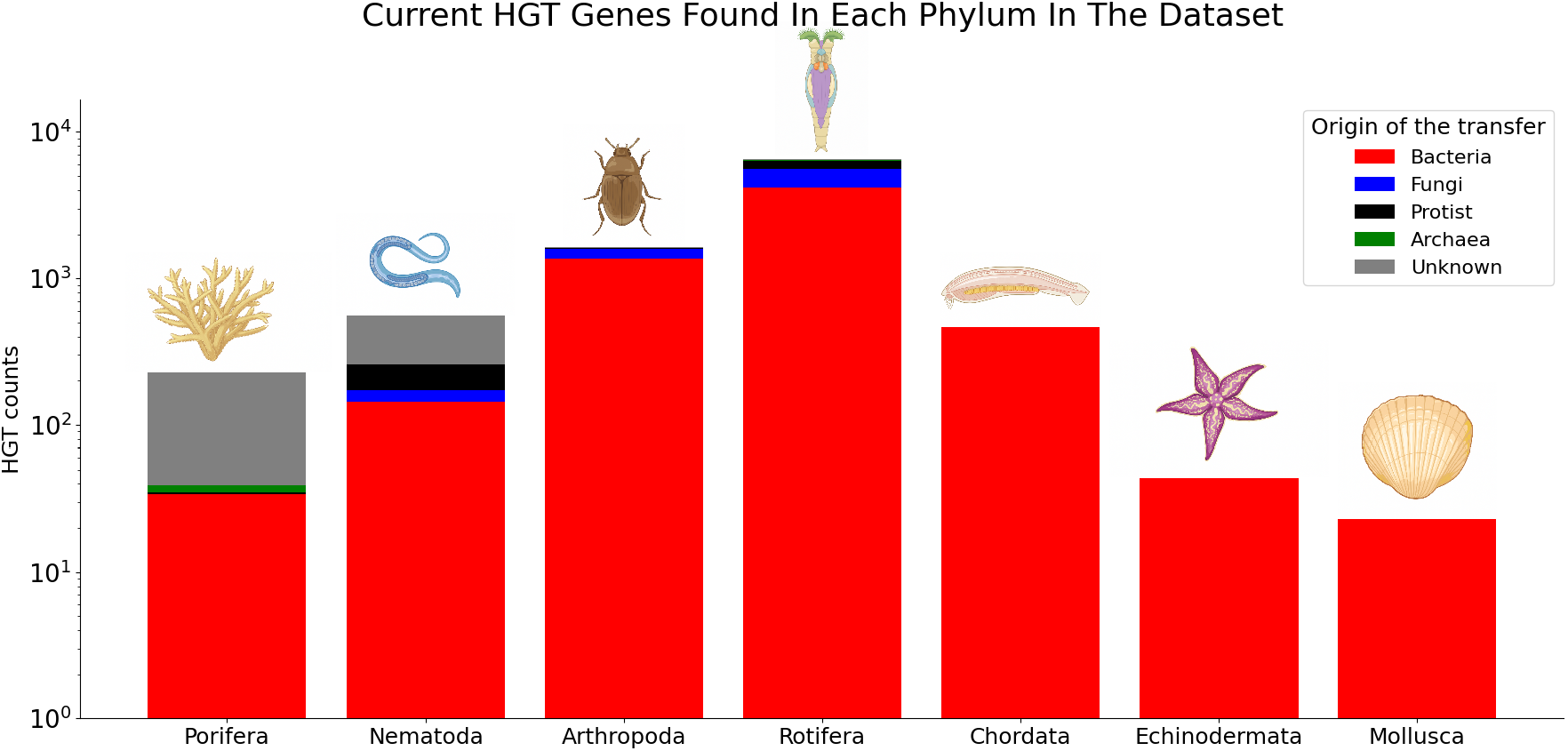
The number of HGTs found in each of the top 7 phyla in the euHGT dataset. ‘Unknown’ are genes with origin not explicitly stated in the article.

### Recurrent phyla that experienced HGT

In total 13 phyla were represented in the euHGT dataset. The phyla containing the most bacterial gene transfers were Rotifera, Arthopoda, Chordata, and Nematoda respectively (Figure 3). Genes with fungal origins were abundant in Rotifera and Arthopoda. Moreover, Rotifera and Nematoda contained the most genes with protist origins. We also assessed the distribution of GC content for the transferred genes in each of the 13 phyla. The majority of the genes were above 20% GC content with the highest GC content gene in Arthopoda (Figure 4).

**Figure 3.**
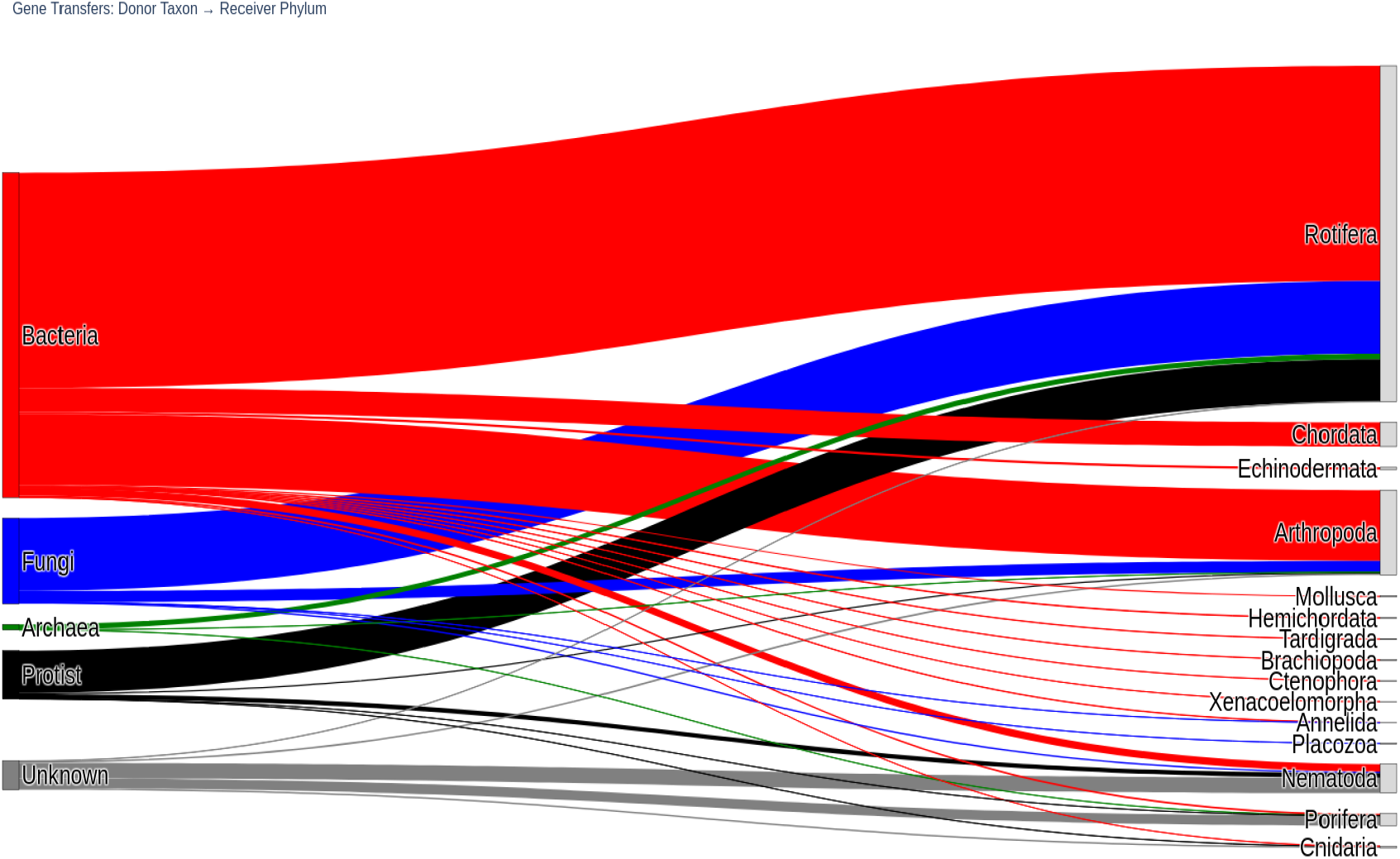
Sankey diagram showing the flow of gene transfers from donor organism to the 13 phyla present in the dataset.

**Figure 4.**
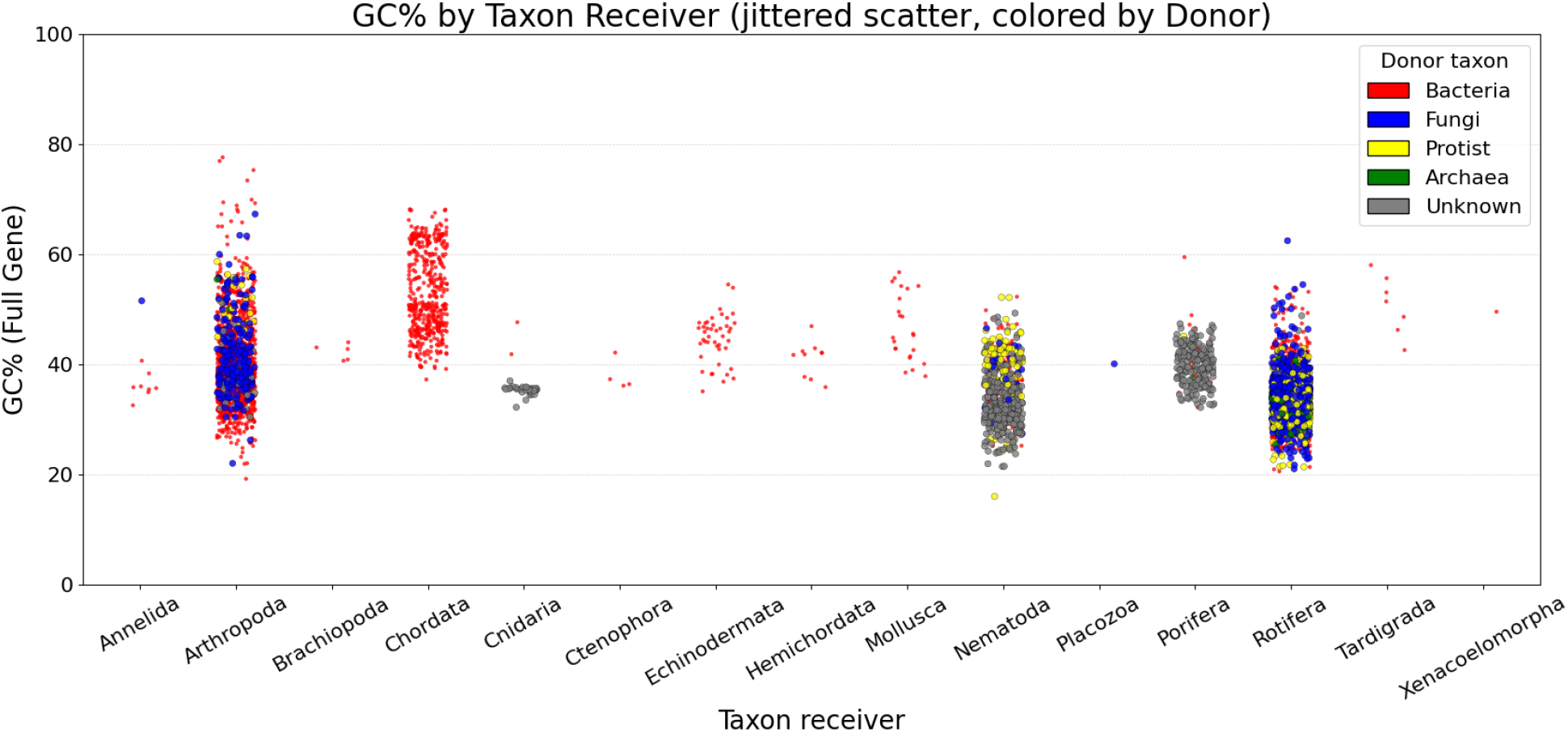
The GC (as % of the gene sequence) for all transferred genes in each phyla. The data are plotted with a slight displacement (jitter) to enhance visualization.

### Data Records

The euHGT dataset is available at: https://figshare.com/articles/journal_contribution/_b_Eukaryotic_horizontal_gene_transfer_dataset_a_compendium_b_/30408499. Lastly, the codes used in this study are available at https://github.com/Genome-explorer/Eukaryotic_horizontal_gene_transfer_dataset_a_compendium. In the dataset each row corresponds to a single gene that was transferred from a donor organism to recipient organism. The columns in the dataset are:

1. Gene_accession_number: The NCBI accession number of the gene or the accession number of the gene in a genome assembly.
2. Genome assembly ID: Represents the genome version used in the paper.
3. Protein_accession_number: The NCBI accession number of the protein or accession number of the protein in a genome assembly.
4. Gene_name/Function: The name of the gene or its function, when available.
5. Full_Gene_Length: The full length of the gene sequence.
6. Coding_Sequence_length (CDS): The length of the coding sequence of the gene.
7. Non_coding_Sequence_length: The number of nucleotides that are not coding for the protein sequence such as 5’UTR, 3’UTR, and intron. This value was calculated by subtracting the CDS length from the full gene length.
8. Amino_Acid_sequence_length (aa): The amino acid sequence length of the protein.
9. G + C_content_full_gene: The decimal fraction of the amount of Guanine-Cytosine in the full gene sequence. The formula: GC Content % = Number of G’s and C’s divided by the total number of bases in the sequence multiplied by 100. If any of the genetic sequences have an “N” in the sequence, the “N” is excluded from the calculation.
10. G + C_content_CDS: The decimal fraction of the amount of Guanine-Cytosine in the CDS gene sequence.
11. Phylogenetic support: Identity’s whether this gene was shown to be transferred using a phylogenetic tree from the paper that identified the gene.
12. level_of_certainty (1-5): A certainty metric used to assess the likelihood of a gene transfer.
13. Taxon_Donor (Bacteria, protist, fungi): The taxon of the Donor organism of the gene that was transferred.
14. Taxon_receiver (By_Phylum): The taxon of the recipient organism that received the gene.
15. Receiver_Genus_species: The organism that received the transferred gene.
16. PMCID/PMID/DOI: The unique paper identifier that is associated with the paper that presented the transferred gene. Primarily the PMCID identifier will be used for each paper, and if a paper does not have a PMCID, then its PMID will be used. Lastly the DOI will be used if both PMCID and PMID are missing.
17. Year: The year the paper was published.
18. Paper: The title of the paper that presented the transferred the gene.

### Technical Validation

The papers that were used to create this dataset have been peer reviewed and published in their respective journals.

## Code availability

Code for the Python scripts that were used to calculate the Guanine-Cytosine percentage, protein length, and coding sequence length for each of the transferred genes is available at https://github.com/Genome-explorer/Eukaryotic_horizontal_gene_transfer_dataset_a_compendium. The code for the web scraper is available at https://github.com/rafsanlab/ScrapPaper.

## Acknowledgments

The authors thank the Fierst lab for support, useful critiques, and suggestions. This research was supported by NSF award 2225796, NSF Louis Stokes Alliances for Minority Participation (LSAMP) Bridge to the Doctorate fellowship award 2204743, and NIGMS award R35GM147245.

## Author information

### Contributions

Johnathan Spaulding collected each of the genes that were identified in the research papers and created the dataset. He also documented the workflow of how to get the genes from each paper and any peculiarities in the process. The custom Python scripts that were used to calculate the Guanine-Cytosine percentage, protein length, coding sequence length, non-coding sequence length, for each of the transferred genes were created by him. Janna Fierst contributed to the dataset design, the refinement process of the figures used in the study, and the writing of the paper.

### Competing interests

The authors declare no competing interests.

All authors declare that there are no financial/commercial conflicts of interest. All copyright permissions have been obtained by the authors.

## Notes

https://figshare.com/articles/journal_contribution/_b_Eukaryotic_horizontal_gene_transfer_dataset_a_compendium_b_/30408499

### Competing Interest Statement

The authors have declared no competing interest.

